# The unique role of parietal cortex in action observation: Functional organization for communicative and manipulative actions

**DOI:** 10.1101/2021.01.22.427829

**Authors:** Burcu A. Urgen, Guy A. Orban

**Affiliations:** Department of Psychology, Bilkent University, 06800, Bilkent, Ankara, Turkey; Interdisciplinary Neuroscience Program, Bilkent University, 06800, Bilkent, Ankara, Turkey; National Magnetic Resonance Research Center (UMRAM) and Aysel Sabuncu Brain Research Center, Bilkent University, 06800, Bilkent, Ankara, Turkey; Department of Medicine and Surgery, Neuroscience Unit, University of Parma, Italy

## Abstract

Action observation is supported by a network of regions in occipito-temporal, parietal, and premotor cortex in primates. Recent research suggests that the parietal node has regions dedicated to different action classes including manipulation, interpersonal, skin-displacing, locomotion, and climbing. The goals of the current study consist of: 1) extending this work with new classes of actions that are communicative and specific to humans, 2) investigating how parietal cortex differs from the occipito-temporal and premotor cortex in representing action classes. Human subjects underwent fMRI scanning while observing three action classes: indirect communication, direct communication, and manipulation, plus two types of control stimuli, static controls which were static frames from the video clips, and dynamic controls consisting of temporally-scrambled optic flow information. Using univariate analysis, MVPA, and representational similarity analysis, our study presents several novel findings. First, we provide further evidence for the anatomical segregation in parietal cortex of different action classes: We have found a new region that is specific for representing human-specific indirect communicative actions in cytoarchitectonic parietal area PFt. Second, we found that the discriminability between action classes was higher in parietal cortex than the other two levels suggesting the coding of action identity information at this level. Finally, our results advocate the use of the control stimuli not just for univariate analysis of complex action videos but also when using multivariate techniques.

## 1. INTRODUCTION

Over the last two decades, neurophysiological and neuroimaging studies in human and nonhuman primates have identified a network of brain regions in occipito-temporal, parietal and premotor cortex that are associated with observing others’ actions, known as the Action Observation Network (AON) (Cross et al 2009a, Caspers et al, 2010; Nelissen et al, 2011; Jastorff et al 2010). One prominent finding that has guided action observation research to date is that it shares some neural resources with action execution including some regions in the parietal and premotor cortex (Rizzolatti and Craighero, 2004). Comparative functional neuroanatomical studies of action execution have demonstrated that posterior parietal cortex (PPC) is functionally divided into parts for different actions including grasping, reaching, hand-to-mouth behavior, looking, and protecting the face or body (Kaas and Stepniewska, 2016; Graziono and Aflalo, 2007). These subdivisions are thought to form “intentional maps” generating preliminary plans for movement (Anderson and Buneo, 2002). The relative expansion of the PPC at the level of the inferior parietal lobule (IPL, Van Essen and Dierker, 2007) in humans suggests that it may host additional maps devoted to actions specific to humans (Orban, 2016; Kaas and Stepniewska, 2016). One implication of this research, when combined with the findings of the shared common neural resources for action execution and observation is that parietal cortex may be functionally organized (Goodale and Milner, 1992) and divided into functionally distinct regions for different observed actions in action observation.

There is a growing body of research in action observation that supports the hypothesis of an anatomical segregation in parietal cortex for different action classes, including manipulation, interpersonal, skin-displacing, locomotion, and climbing (Jastorff et al, 2010, Abdollahi et al 2013, Ferri et al, 2015). The primary aim of the current study is to build upon this work with two new classes of actions that are communicative in nature, at least one of which is specific to humans (Liebal and Call, 2012, Graham et al 2018). More specifically we are interested in whether there are parietal regions that are specifically activated by direct and indirect communicative hand actions, and if so where they are localized in the IPL of the human brain. Direct communicative actions are symbolic hand actions addressed to a conspecific present in the scene, while indirect communication actions involve leaving a symbolic trace (shape) in a substrate that can be viewed later by a conspecific.

A second aim of the current study is to investigate how the three levels of the AON differ from one another in representing action classes. Computational modeling (Giese and Poggio, 2003) and single cell studies in monkeys (Vangeneugden et al 2009) suggest that form and motion, both components of an observed action, are processed in the ventral and dorsal pathways of the visual system respectively, and that this information is sent to the occipito-temporal cortex, i.e. the first level of the AON, to be integrated. This visual information is passed on to the other two levels of the AON, parietal and premotor cortex for further processing (Nelissen et al 2011). What kind of computations are being carried out in these levels and how they differ from each other is not well understood (Cross et al, 2009b, Wurm et al, 2015, Chan and Baker, 2015; Urgen et al, 2019). The present study also aims to shed light on this issue with a particular focus on how observed action classes are represented at the three levels of the AON.

To this end, we used both univariate and multivariate techniques in a complementary fashion. Using a whole-brain approach and univariate analysis, we functionally localized the regions activated by three different classes of actions as well as the common and specific regions for each action class (Abdollahi et al 2013). Next, using multivariate pattern analysis (MVPA) (Pereira et al 2009) we sought to identify which levels of the AON could discriminate between the action classes, and how they differed. Finally, using representational similarity analysis (RSA) (Kriegeskorte et al 2008, Liu et al 2013), we compared the neural representational distances between the action classes and their exemplars across ROIs to reveal the functional properties at each level of the AON.

## 2. METHODS

### 2.1 Participants

32 subjects (15 females, 17 males, mean age = 26, SD = 2.87) participated in the study. All participants were right-handed, had normal or corrected-to-normal visual acuity and no history of mental illness or neurological diseases. The study was approved by the Ethical Committee of Parma Province and all volunteers gave written informed consent in accordance with the Helsinki Declaration prior to the experiment.

### 2.2 Stimuli

The stimuli of the experiment consisted of short video clips (448 x 336, 50 fps, 2.6 seconds) showing an actor from the left side while performing three classes of actions: (1) Indirect communication (Figure 1A), (2) Direct communication (Figure 1B), and (3) Manipulation (Figure 1C).

**Figure 1.**
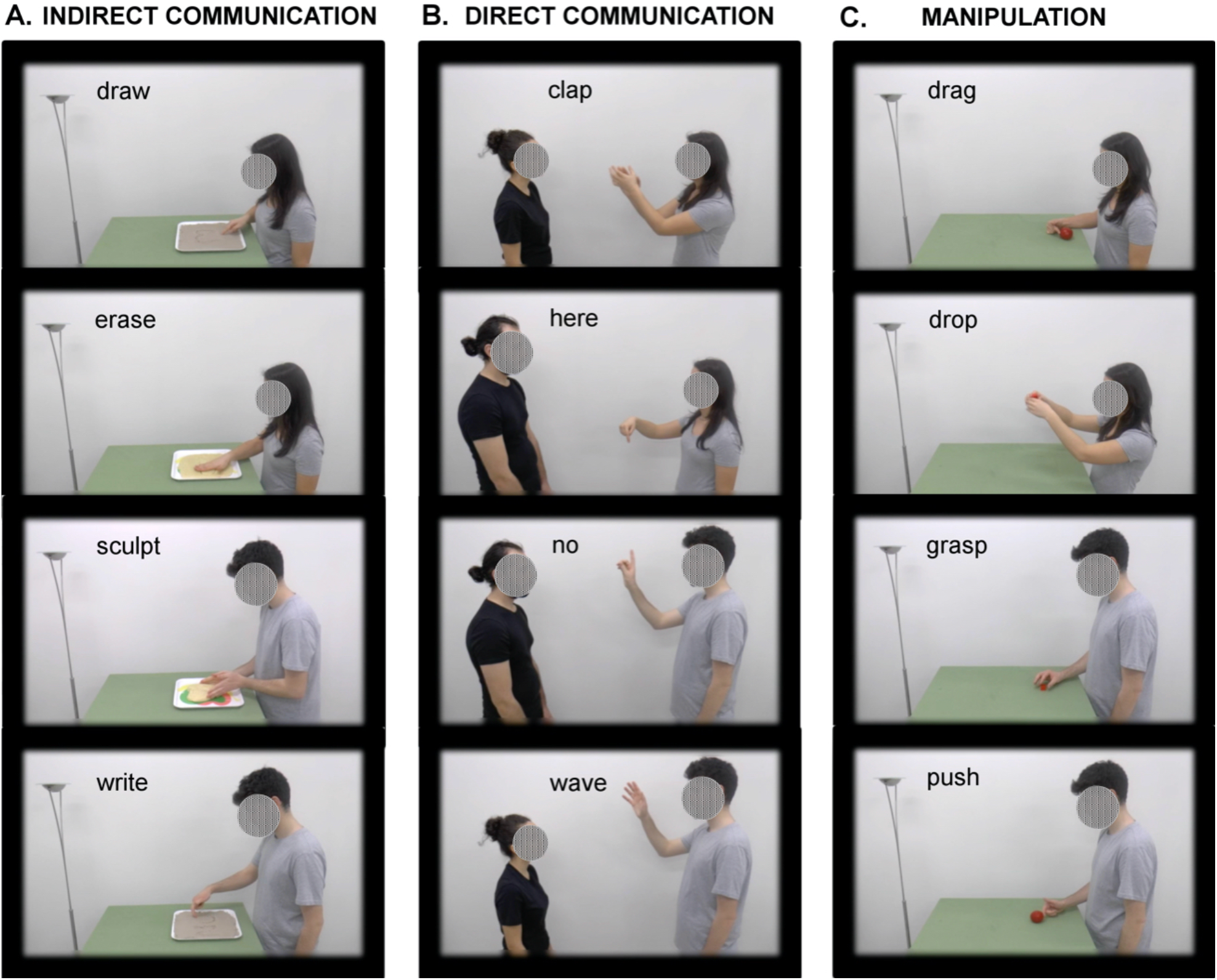
Static frames from the videos of the three action classes and their four exemplars in the experiment. (A) Indirect communication, (B) Direct communication, (C) Manipulation.

All actions included finger or hand movements. Each *action class* included four *exemplars*, and each exemplar had four *variants*, yielding 16 different video clips per action class. The exemplars of the indirect communication class included writing with the index finger, drawing with the index finger, erasing with the right hand, and sculpting with both hands. All of these were performed on two different types of substrate, sand or dough, by a male or a female actor (which made up the four variants for a given exemplar) to communicate a message (write ‘no’, draw a heart, erase ‘Ciao’ (‘Hello’ in Italian), or sculpt a heart). The exemplars of the direct communication class included waving with the right hand, clapping with both hands, displaying “no” sign by moving the index finger left and right continuously, and displaying “right here” by moving the index finger up and down continuously. Each of these actions was performed by a male or a female actor, and was targeted toward a male or female actor (which made up the four variants for a given exemplar). The exemplars of the manipulation class included grasping an object with the right hand, dropping an object with both hands, dragging an object toward the body with the index finger, and pushing an object away from the body with the index finger. The exemplars are the same as in previous work (Ferri et al 2015), but the effectors were slightly changed to match those of communicative actions. Each of the manipulative actions were targeted toward a small or a large object, and performed by a male or female actor (which made up the four variants for a given exemplar). We matched the action classes to each other as much as possible (see Table 1 for stimuli properties) by including the same two actors with the same clothing, a target for the action, and equal number of actions performed by a single finger, single hand (unimanual) or two hands (bimanual), and same number of elements in the scene.

In addition to the action stimuli (48 videos in total), two different control stimuli were used: static controls (SC), which consisted of the first, middle, and last frames of each action video, and dynamic controls (DC), which consisted of the optical flow extracted from each action video, superimposed onto a noise pattern, temporally de-correlated between squares of a superimposed grid and reduced to average translation within a square (similar to Ferri et al 2015). The rationale was to control for the two components of an observed action: a figural component (shape of the body and scene), and a motion component (motion vectors of the body). Finally, in all videos or static frames, a blue fixation dot was present near the position where the hand action unfolded. In half of the videos and static frames the fixation dot was slightly above the middle of the effector trajectory, and in the other half, slightly below the effector. Each video or frame was positioned on the screen in such a way that the fixation dot always remained at the center.

**Table 1.** Properties of the action video stimuli

| Table 1. Properties of the action video stimuli |  |  |  |  |
| --- | --- | --- | --- | --- |
| ACTION CLASS/<br>EXEMPLARS | ACTOR | TARGET | MANUAL/<br>BIMANUAL | FINGERS |
| **INDIRECT**<br>**COMMUNICATION** |  |  |  |  |
| Write | Male/Female | Sand/Dough | Manual | Single |
| Draw | Male/Female | Sand/Dough | Manual | Single |
| Erase | Male/Female | Sand/Dough | Manual | All five |
| Sculpt | Male/Female | Sand/Dough | Bimanual | All ten |
| **DIRECT**<br>**COMMUNICATION** |  |  |  |  |
| “Here” gesture | Male/Female | Male/Female | Manual | Single |
| “No” gesture | Male/Female | Male/Female | Manual | Single |
| Wave | Male/Female | Male/Female | Manual | All five |
| Clap | Male/Female | Male/Female | Bimanual | All ten |
| **MANIPULATION** |  |  |  |  |
| Drag | Male/Female | Big/Small | Manual | Single |
| Push | Male/Female | Big/Small | Manual | Single |
| Grasp | Male/Female | Big/Small | Manual | All five |
| Drop | Male/Female | Big/Small | Bimanual | All ten |

### 2.3 Experimental Design

The experiment consisted of 10 runs following a block design. In each run, the three action classes (3 blocks), and one of the controls (SC or DC) corresponding to the three action classes (3 blocks) were presented in a randomized order, and followed by a fixation block lasting as long as the stimulus blocks (41.6 seconds). This was repeated twice in each run, lasting 582.4s (14×41.6s). SC and DC were presented in alternating runs. In each SC run, one type of static frame was used (first, middle, or last frame), with their order randomized across SC runs. Within each block 16 videos/images of any given class (4 exemplars x 4 variants for a given exemplar) were presented for a total of 41.6s (16×2.6s). Within a block the 4 exemplars of a given class were presented in a randomized order, but the 4 variants of each exemplar were presented one after the other, defining mini-blocks of exemplars within a given block. The order of the variants was fully randomized as well.

### 2.4 Data Collection and Data Preprocessing

We scanned our subjects at the University Hospital of University of Parma using 3T MR scanner (GE Discovery MR750, Milwaukee, ILL) with an 8-parallel channels receiver coil. Subjects were presented the stimuli visually through a head-mounted display (Resonance Technology, Northridge, CA, 60 Hz refresh rate) with a resolution of 800 x 600 pixels while they laid in the scanner, and were asked to fixate a dot at the center of the screen throughout the experiment. The presentation of the stimuli was controlled by E-prime software (Psychology Software Tools, Sharpsburg, PA). In order to reduce head motion during scanning, subjects’ heads were supported with cushion pads. During the presentation of the stimuli, eye movements were recorded with an infrared eye tracking system (60 Hz, Resonance Technology, Northridge, CA).

We acquired functional images during the presentation of the stimuli followed by the acquisition of a high-resolution structural image for anatomical reference. The functional images were acquired using gradient-echoplanar imaging with the following parameters: TR = 3 seconds, TE = 30 ms, flip angle = 90, 96 x 96 matrix with FOV 240 (2.5 x 2.5 mm in plane resolution), 49 horizontal slices (2.5 mm thickness, and 0.25 mm gap), ASSET factor of 2. The 49 slices covered the whole brain including cerebellum. In total, 1950 volumes (195 x 10 runs) were collected in the experimental session. The structural image was a T1-weighted IR-prepared fast SPGR image and covered the whole brain with 186 sagittal slices with 1 x 1 x 1 mm^3^ resolution. Its acquisition parameters were as follows: TE/TR 3.7/9.2 ms, inversion time = 650 ms, flip angle = 12, acceleration factor (ARC) = 2.

### 2.5 Data Preprocessing and Univariate analysis

Data preprocessing and univariate analysis were performed using SPM8 software package (http://www.fil.ion.ucl.ac.uk/spm/software/spm8/) running under MATLAB (The Mathworks, Natick, MA). The fMRI data of each subject were pre-processed with standard procedures including motion correction, coregistration of the anatomical image and the mean functional image, spatial normalization of all images to a standard stereotaxic space (MNI) with a voxel size of 2 x 2 x 2 mm, and smoothing of the resulting images with Gaussian kernel of 6 mm. We discarded 5 subjects who did not fixate well according to the eye movement data (i.e. they made more than 15 saccades per minute for any of the conditions). We thus included 27 subjects (13 females, 14 males) in the final analysis. These subjects averaged 8.4 saccades per minute. Although there was a main effect of condition when we performed a one-way ANOVA on the saccades data considering all experimental conditions and fixation (F_6,156_ = 9.51, p<0.01), the effect was not driven by the differences between the experimental conditions but rather primarily by the larger number of saccades in the fixation condition (9.7 on average) compared to the experimental conditions since pair-wise comparisons between the experimental conditions were not significant (p>0.1). Thus any difference in the brain activity patterns between the experimental conditions cannot be explained by differences in fixation.

After preprocessing, we performed a first-level analysis for each subject using a General Linear Model (GLM) with condition onsets and durations. The design matrix consisted of 13 regressors including 7 experimental conditions (3 action classes, 3 controls (DC or SC), and 1 fixation), and 6 motion parameters (3 rotations, 3 translations). All regressors were convolved with the canonical hemodynamic response function. Next, we calculated contrast images on the SPM generated in the first level, and then performed a second-level random effects analysis (Holmes and Friston, 1998) for the 27 subjects included in the analysis. We defined three types of statistical maps: Activation map, Specific map, and Common activation map (see below). For these maps, the simple or interaction contrasts were defined at the first level, while conjunctions and masking were made at the second level.

#### 2.5.1 Whole-brain Analysis: Activation Maps

We defined various contrasts in order to identify statistical activation maps for each action class (similar to Ferri et al 2015*). Activation maps* correspond to the network of brain regions that were significantly more strongly activated by the observation of a particular action class than the observation of the static or dynamic controls of that action class. The activation map for each action class was defined by the contrast [Action – (0.5*DC + 0.5*SC)] and masked inclusively by [Action - Fixation] at p<0.01. Please note that DC and SC in each contrast refer to the dynamic and static controls of the respective action. We included in the activation map for each action class those clusters that survived the p<0.05 FWE (corrected) at peak level. The activations maps defined in SPM8 were projected onto the flat surface of the left and right hemispheres of the human PALS B12 atlas (Van Essen, 2005) using the Caret software package (Van Essen et al, 2001) since flat maps allow better visualization of sulci and gyri as well as displaying all three views (axial, sagittal, and coronal) of the hemisphere in a single image.

#### 2.5.2 Whole-brain Analysis: Specific Maps and Common Activation Maps

We defined *specific maps* in order to identify the regions that were activated solely by observing a particular action class (similar to Ferri et al 2015). The specific map for each action class was defined by the conjunction of the two contrasts 1) [(Action1 - (0.5*DC + 0.5*SC)) – (Action2 - (0.5*DC + 0.5*SC)], 2) [(Action1 - (0.5*DC + 0.5*SC)) – (Action3 - (0.5*DC + 0.5*SC)], masked inclusively by the activation map of the action class of interest (Action1) at p<0.01, and the contrast [Action1 - Fixation] at p<0.01, and exclusively by the activation maps of the other two action classes also at p<0.01.

We also defined the common activation map across observed action classes, which correspond to the network of brain regions that were significantly activated by all three action classes compared to their static and dynamic controls. As in previous studies (Abdollahi et al 2013, Ferri et al 2015), these were defined by the conjunction of three contrasts 1) [Action1 - (0.5*DC + 0.5*SC)], 2) [(Action2 - (0.5*DC + 0.5*SC))], and 3) [Action3 - (0.5*DC + 0.5*SC)] with a conjunction threshold of p<0.001 (uncorrected). The specific and common activation maps defined in SPM8 were then projected onto the left and right hemispheres of the human PALS B12 atlas (Van Essen, 2005) using the Caret software package (Van Essen et al, 2001).

#### 2.5.3 Activity Profiles

We defined the *activity profiles* as the mean percent signal change in the BOLD response for a given condition compared to the baseline (fixation condition) in a region of interest (ROI). We computed the activity profiles for *action classes* for the three levels of the Action Observation Network using eight a priori defined ROIs (2 ROIs in occipito-temporal, 5 ROIs in parietal, and 1 ROI in premotor cortex, see the section *2.6* *Identification of ROIs*).

Next, we performed an 8 (ROI) x 2 (Hemisphere) x 3 (Action class) x 3 (Presentation mode) mixed ANOVA on the activity profiles for statistical analysis, the four factors being ROI (OTS, MTG, DIPSM, DIPSA, phAIP, PFt, PFcm, Premotor), Hemisphere (Left, Right), Action classes (Indirect communication, Direct communication, Manipulation), and Presentation mode (Video, Static control, Dynamic control).

### 2.6 Identification of ROIs

We defined two sets of independent ROIs to cover the three levels of the Action Observation Network bilaterally (Figure 2) using functional or cytoarchitectonic criteria.

**Figure 2.**
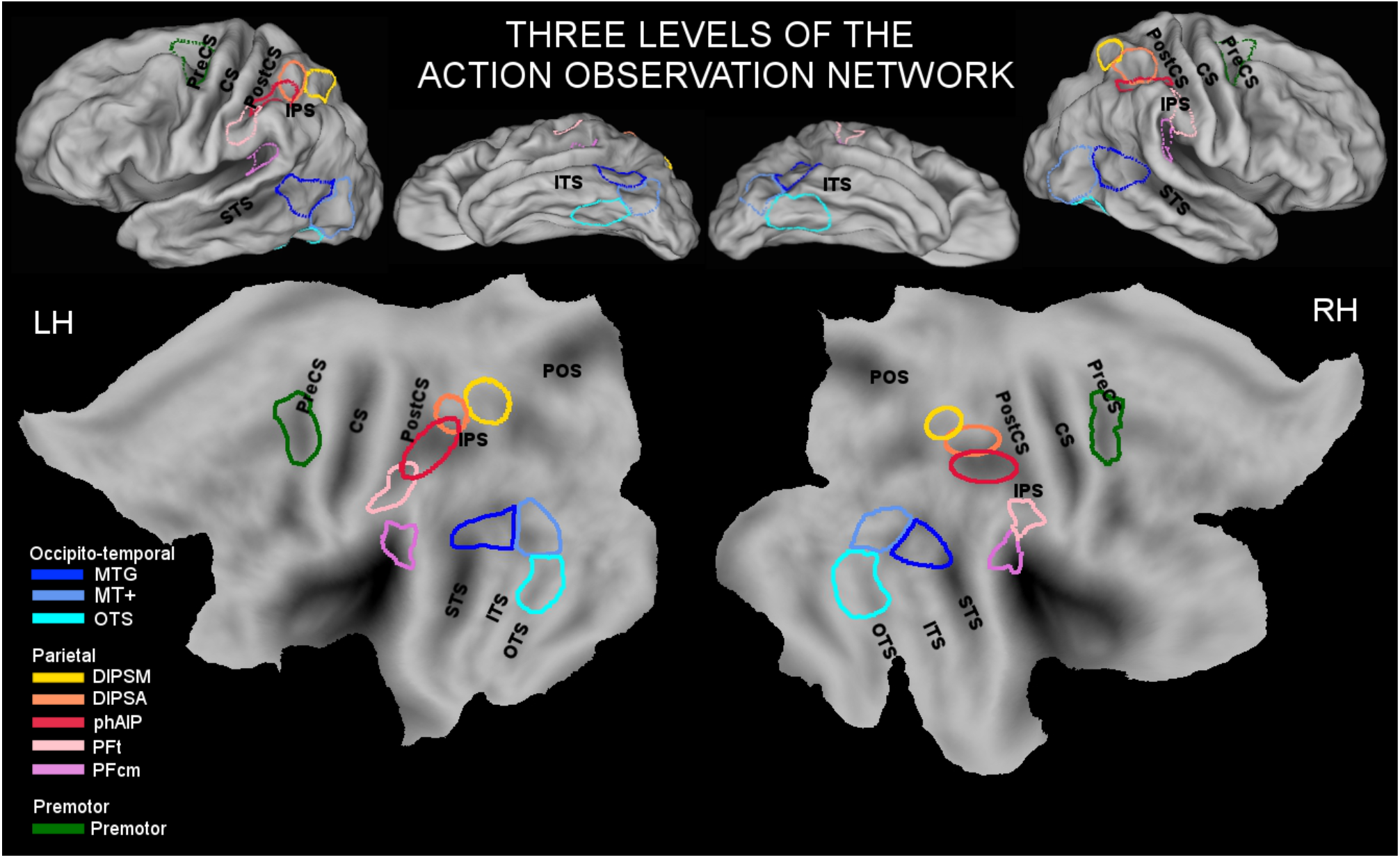
ROIs included in the three levels of the Action Observation Network (occipito-temporal, parietal, and premotor) displayed on surface maps (upper panel) and flat maps (lower panel) for the left (LH) and right hemispheres (RH). Anatomical landmarks are marked in black: CS - Central sulcus, PreCS – Precentral sulcus, PostCS – Postcentral sulcus, IPS – Intraparietal sulcus, POS – Parieto-occipital sulcus, STS – Superior temporal culcus, ITS – Inferior temporal sulcus, OTS – Occipito-temporal sulcus.

The functional ROIs included middle temporal gyrus (MTG) and occipito-temporal sulcus (OTS) in the occipito-temporal cortex, and premotor cortex (PMC) taken from the maps of Ferri et al (2015), plus the confidence ellipses dorsal intraparietal sulcus medial (DIPSM), dorsal intraparietal sulcus anterior (DIPSA), and putative human AIP (phAIP) in the parietal cortex (Jastorff et al. 2010). The phAIP and DIPSA (more precisely its ventral 2/3) regions correspond to the rostral motor and caudal visual parts of monkey anterior intraparietal (AIP) area, and DIPSM to anterior part of the lateral intraparietal (LIP) area of the monkey (Orban 2016). The cytoarchitectonic ROIs included PFt and PFcm in the inferior parietal lobule (Caspers et al. 2006).

### 2.7 Multivariate Techniques: MVPA and Representational Similarity Analysis

#### 2.7.1 MVPA

We performed a 3-way classification between action classes (Indirect Communication, Direct Communication and Manipulation) on the brain activity patterns for each presentation mode, i.e. videos, static controls, and dynamic controls in all ROIs (MTG, OTS, DIPSM, DIPSA, phAIP, PFt, PFcm, and PMC bilaterally).

To prepare the data for classification, we first performed another GLM. In this GLM analysis, each exemplar trial was treated as a different condition regardless of the variants. In other words, the BOLD response was convolved with the mini-blocks instead of the main block (See section 2.5 Univariate Analysis above). The beta images generated in the first level were fed into the classifier.

In order to perform the classifications, we used support vector machines (SVM) (Cortes and Vapnik, 1995) with a linear basis function and LIBSVM software package (Chang and Lin, 2011). The data were separated into a training set (80% of the total data), and test set (20% of the total data), and both were scaled before classification. Five-fold cross-validation scheme (Pereira et al. 2009) was applied during training.

The prediction accuracy (the ratio of the number of correct predictions to all predictions) was used as the performance metric of the classifier. For the 3-way between-class classifications, the upper limit of above-chance performance with 95% confidence interval was 41% in videos, and 44% in static controls and dynamic controls, computed using the algorithm of Muller-Putz et al (2008).

After computing the prediction accuracy for each ROI, we performed an 8 (ROI) x 2 (Hemisphere) x 3 (Presentation mode) ANOVA to compare the discriminability of actions across ROIs and presentation modes for 3-way prediction accuracies.

#### 2.7.2 Representational Similarity Analysis (RSA)

In order to study the functional properties of the brain regions involved in action processing, and to investigate the representational space of the brain regions for actions, we used representational similarity analysis (Kriegeskorte et al, 2008). This technique allows investigators to quantify how similar the neural patterns are that correspond to different conditions of an experiment within a certain brain region. In addition, the application of clustering methods (e.g. hierarchical clustering or multidimensional scaling) to the similarity measures allows one to study the representational space of the particular brain region. Further, comparison of the similarity structures across brain regions allows one to study how neural representations evolve along the cortical hierarchy.

We calculated the representational dissimilarity matrix (RDM) in each region of interest (ROI) for each subject by taking the correlation distance (1 - correlation) between all pairs for action exemplars using the beta images derived in the first-level (univariate) analysis in SPM8, which resulted in a 12 x 12 matrix (3 action classes x 4 exemplars for each action class). This was done for the action videos as well as for the static and dynamic controls. For each condition, we computed the grand average dissimilarity matrix by taking the average of all subjects for each ROI. We then applied hierarchical clustering on the grand average RDMs to examine the similarity patterns for action classes, and used multidimensional scaling (MDS) to visualize the geometric distance between action classes. After MDS was applied to the dissimilarity matrices, we computed the average distance of each action class to the other two action classes: d_ij_ = MDS distance between (Action class)_i_ and (Action class)_j_, and summed them up to compute a distance metric for each ROI. All the steps in RSA were performed using custom scripts in MATLAB.

## 3. RESULTS

### 3.1 Univariate Analysis

#### 3.1.1 Activation Maps for Action Classes

The first step in the univariate analysis was to functionally localize regions involved in the observation of the three action classes, indirect communication, direct communication, and manipulation. These regions were shown in the activation map of each action class (Figure 3). All three maps were bilateral, but showed a clear bias favoring the left hemisphere.

**Figure 3.**
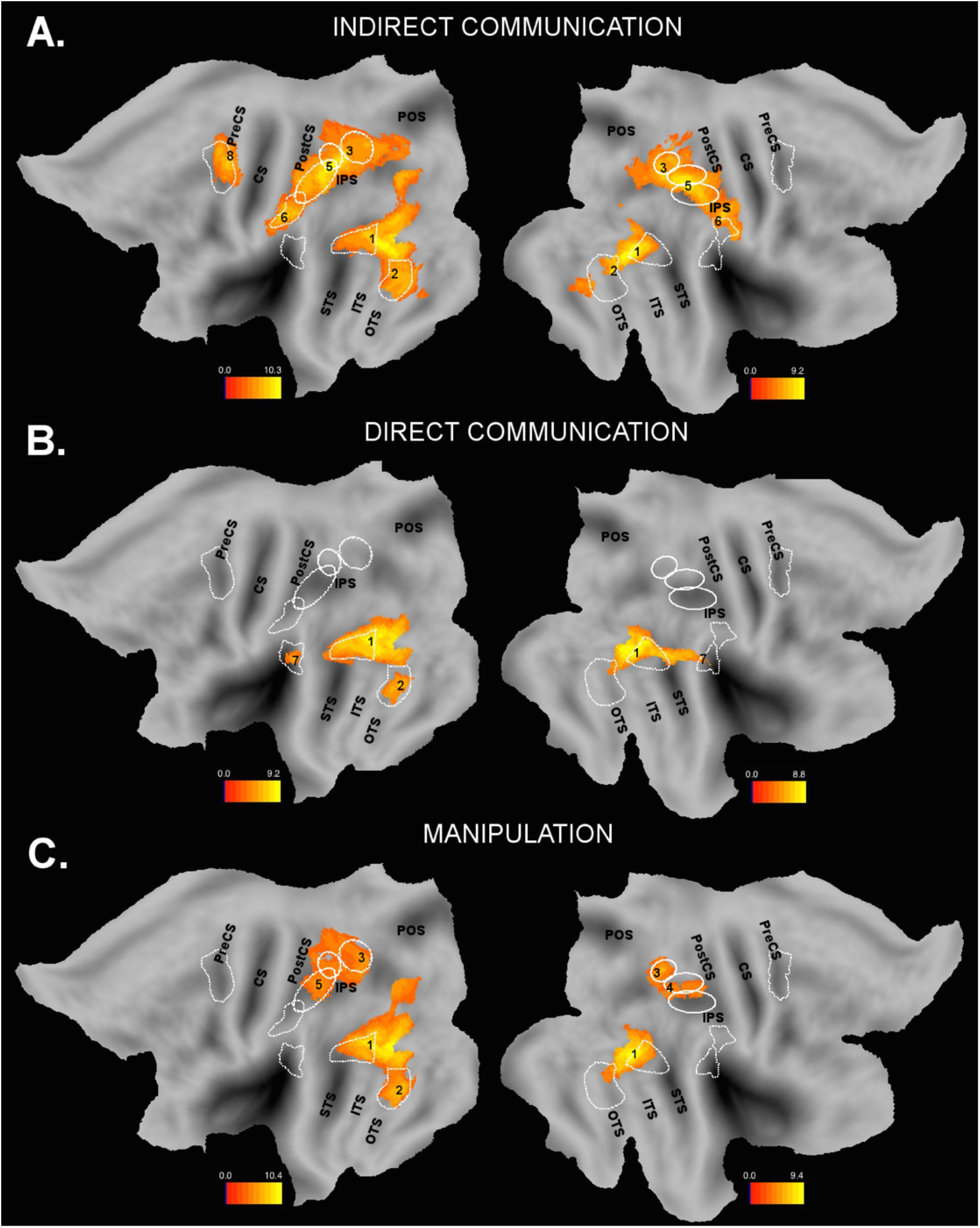
Activation maps for each action class. The white lines indicate the borders of the ROIs included in the study as shown in Figure 2. The anatomical landmarks are marked as in Figure 2. (A) Activation map for Indirect communication (p<0.05 FWE corrected at peak level). Local maxima were labeled with numbers whose coordinates were listed in Table 2. Color bar shows the t-values. (B) Activation map for Direct communication (p<0.05 FWE corrected at peak level). Local maxima were labeled with numbers whose coordinates were listed in Table 2. (C) Activation map for Manipulation (p<0.05 FWE corrected at peak level). Local maxima were labeled with numbers whose coordinates were listed in Table 2.

The activation map for indirect communication included areas at all three levels of the Action Observation Network (Jastorff et al 2010): MTG and OTS as well as the MT+ bilaterally in the occipito-temporal cortex (OTC); DIPSM, DIPSA, phAIP, PFt bilaterally in parietal cortex, and left premotor cortex (Figure 3A). Although the activation pattern was bilateral, it was left biased especially at the OTC and PMC levels. The local maxima of the activations in each ROI are labeled with numbers on the flat map, and the corresponding SPM coordinates shown in Table 2 with the corresponding numbers. The local maxima in MTG (bilateral) were located in the posterior portion, near the border with MT+, whereas the local maxima in OTS (bilateral) were in the upper part. The local maxima in DIPSM (bilateral) were near the border with DIPSA, as were those in phAIP (bilateral). For PFt, the local maximum was located in its rostral and caudal parts in the left and right hemisphere, respectively. The local maxima in the left premotor cortex was in the posterior part.

**Table 2.** Local Maxima of the SPM activation maps (p<0.05 FWE at peak level)

| Table 2. Local Maxima of the SPM activation maps ($p < 0.05$ FWE at peak level) | | | | | | |
| --- | --- | --- | --- | --- | --- | --- |
|  | Indirect Communication |  | Direct Communication |  | Manipulation |  |
|  | Left | Right | Left | Right | Left | Right |
| **Occipito-temporal** |  |  |  |  |  |  |
| 1. MTG | -44 -70 4 | 46 -62 4 | -48 -68 6 | 46 -64 6 | -48 -68 4 | 46 -64 6 |
| 2. OTS | -42 -60 -10 | 42 -62 -12 | -39 -58 -18 |  | -39 -58 -19 |  |
| **Parietal** |  |  |  |  |  |  |
| 3. DIPSM | -30 -58 52 | 24 -62 50 |  |  | -26 -64 48 | 22 -63 53 |
| 4. DIPSA |  |  |  |  |  | 29 -55 48 |
| 5. phAIP | -36 -48 48 | 32 -48 46 |  |  | -38 -44 42 |  |
| 6. PFt | -56 -27 32 | 52 -27 44 |  |  |  |  |
| 7. PFcm |  |  | -46 -42 20 | 54 -36 24 |  |  |
| **Premotor** |  |  |  |  |  |  |
| 8. Premotor | -30 -14 58 |  |  |  | -28 -12 52<br>(SVC) |  |

The activation map for direct communication included areas MTG and MT+ bilaterally, and left OTS in the occipito-temporal cortex, and bilateral PFcm in the parietal cortex (Figure 3B). The local maxima in MTG (bilateral) were in the posterior part near the border with MT+, whereas the local maxima of the other sites were located near their centers (See Table 2 for the coordinates).

The activation map for manipulation included left MTG, OTS and MT+ as well as right MTG and MT+ with a small extension into OTS in the occipito-temporal cortex similar to the maps for indirect communication and direct communication. DIPSM, DIPSA, and phAIP were activated bilaterally in the parietal cortex, but with a clear left bias (Figure 3C). There was also a small cluster in left premotor cortex after small volume correction (SVC) using premotor activation site of the manipulation map from Ferri et al (2015) as the a priori ROI (10 mm sphere). The local maxima of its coordinates are listed in Table 2. The local maxima in MTG (bilateral) were in the posterior part near the border with MT+, whereas the local maximum in the left OTS was in the middle of the site. In the parietal cortex, the local maxima of DIPSM (bilateral) were located in its posterior part, of right DIPSA in the posterior portion near the border with DIPSM, and of left phAIP in the center of its posterior half.

#### 3.1.2 Specific Maps and Common Activation Map

After identifying the activation maps for each class, the second step in the univariate analysis was to identify the regions that were *specifically* activated by observing a particular class (specific map), and the regions that were *commonly* activated by observing the three action classes (common activation map).

The specific map for indirect communication was symmetrical, and included a small cluster near OTS bilaterally, the rostral portion of phAIP bilaterally, the rostral portion of PFt bilaterally, a small cluster in the dorsal portion of left premotor, and two small clusters in the dorsal and ventral portions of the right premotor cortex (Figure 4A). The coordinates of the local maxima were shown in Table 3. We did not find any clusters exclusively involved in the observation of direct communication or manipulation actions (i.e. the specific maps for these classes were empty).

**Table 3.**
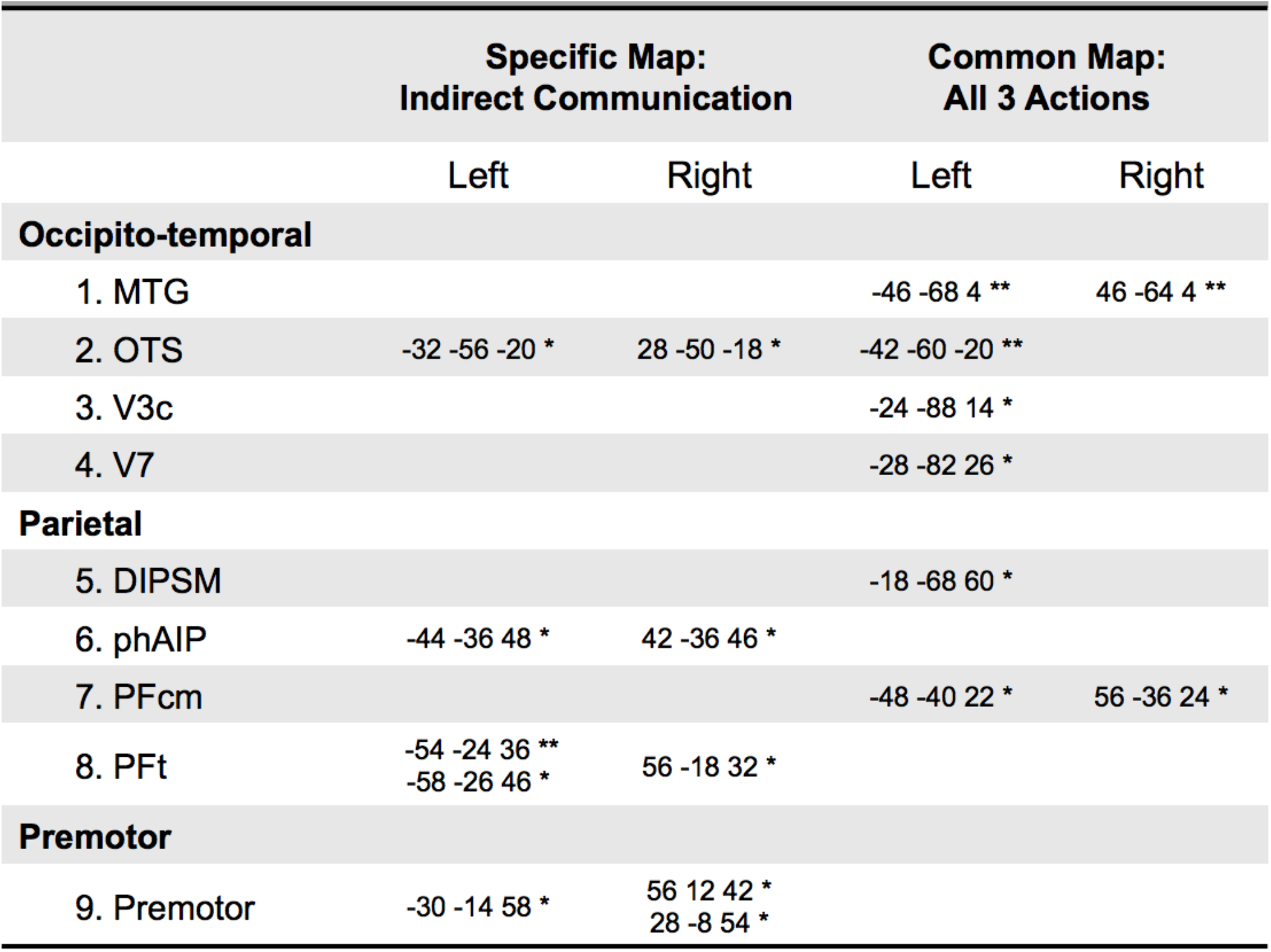
Local Maxima in each ROI of the SPM common map and specific map for indirect communication (*p<0.001 uncorrected; **p<0.05 FWE at peak level)

|  |  |  |  |  |
| --- | --- | --- | --- | --- |
|  | Specific Map:<br>Indirect Communication |  | Common Map:<br>All 3 Actions |  |
|  | Left | Right | Left | Right |
| **Occipito-temporal** |  |  |  |  |
| 1. MTG |  |  | -46 -68 4 ** | 46 -64 4 ** |
| 2. OTS | -32 -56 -20 * | 28 -50 -18 * | -42 -60 -20 ** |  |
| 3. V3c |  |  | -24 -88 14 * |  |
| 4. V7 |  |  | -28 -82 26 * |  |
| **Parietal** |  |  |  |  |
| 5. DIPSM |  |  | -18 -68 60 * |  |
| 6. phAIP | -44 -36 48 * | 42 -36 46 * |  |  |
| 7. PFcm |  |  | -48 -40 22 * | 56 -36 24 * |
| 8. PFt | -54 -24 36 **<br>-58 -26 46 * | 56 -18 32 * |  |  |
| **Premotor** |  |  |  |  |
| 9. Premotor | -30 -14 58 * | 56 12 42 *<br>28 -8 54 * |  |  |

**Figure 4.**
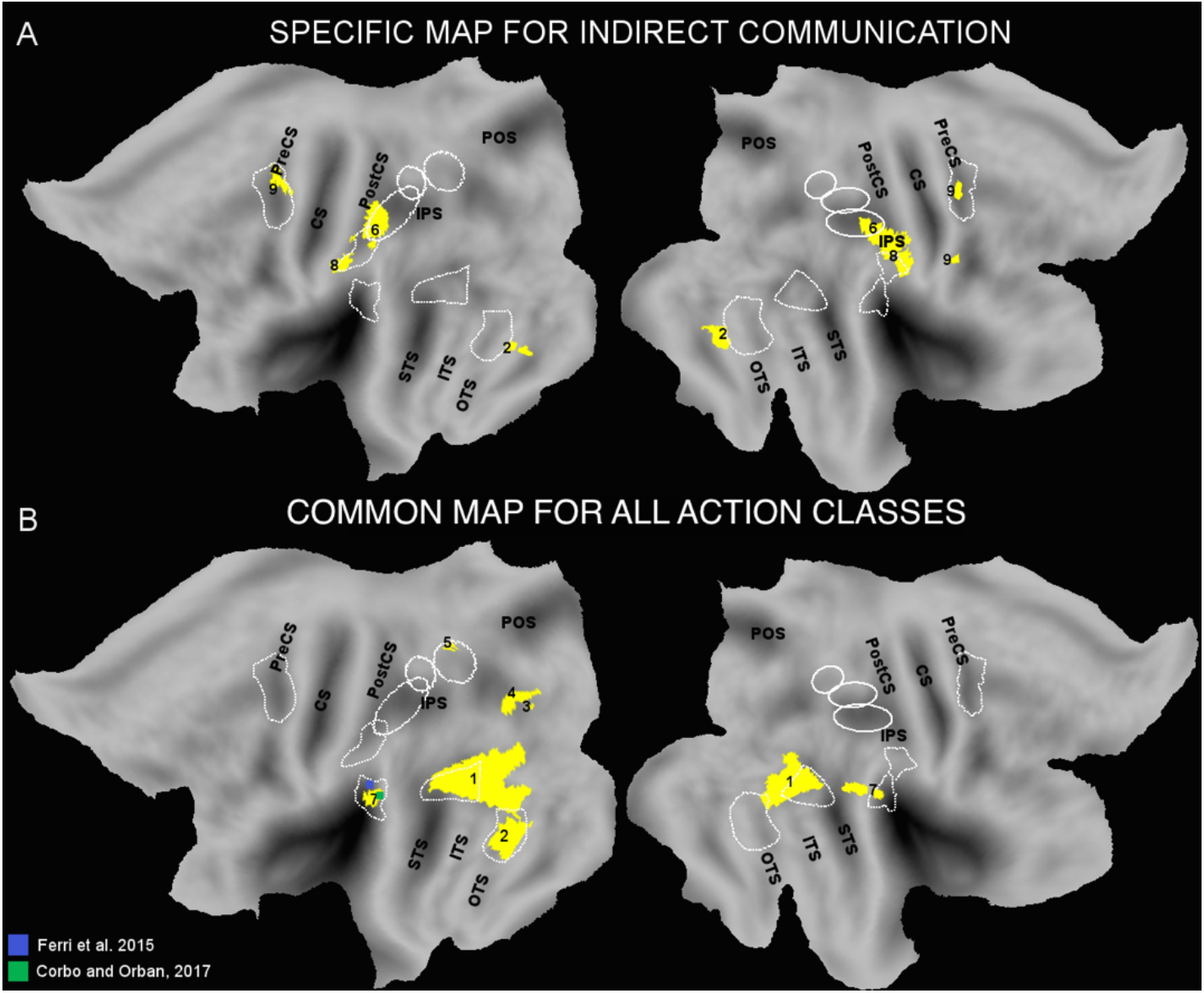
Specific map and Common activation map. (A) Specific map for Indirect communication (p<0.001). The anatomical landmarks are marked as in Figure 2, and the numbers indicate local maxima. (B) Common activation maps of all three action classes, Indirect communication, Direct communication, and Manipulation (p<0.001). The anatomical landmarks are marked as in Figure 2, and the numbers indicate local maxima. The blue and green dots represent the coordinates from previous studies: observed manipulation actions (Ferri et al. 2016) and observed vocal communication actions (Corbo and Orban 2017).

Next, we identified the common activation map for all action classes (Figure 4B). The activation map showed a bias favoring the left hemisphere, as did the activation maps. This map included areas in MTG and MT+ bilaterally and left OTS in occipito-temporal cortex as well as a small cluster in the left visual area V3c (Abdollahi et al 2014). The local maxima in MTG (bilateral) were located in the posterior portion near the border with MT+, whereas the local maximum in the left OTS was in the middle of the site (See Table 3 for the coordinates). There were three common areas in the parietal cortex: PFcm bilaterally, left DIPSM and left V7 (Abdollahi et al 2014). In all parietal sites, the local maxima were located near their centers.

By definition, the specific map shows the cortical regions in which action classes are most differentiated, whereas the common activation map shows the regions in which action classes are most similar in terms of brain activity. The common map for all three classes (Figure 4B) included large activation sites in occipito-temporal cortex but relatively few surface nodes in PPC, while the specific map for indirect communication (Figure 4A) clearly included more nodes in PPC than in occipito-temporal cortex. Indeed, the occipito-temporal activation sites were relatively similar in all 3 activation maps (Figure 3), while these maps showed clear differences at the PPC level. In order to quantitatively compare in which of the three levels of the AON actions were most distinct and most similar, we computed the number of voxels that were active for a given threshold (p<0.001 and p<0.01) around the local maxima of the SPMs of the specific map and the common activation map for the 3 classes. At both thresholds, voxel counts in the parietal cortex outnumbered the other two levels, particularly significantly in the occipito-temporal level, in the specific map, whereas occipito-temporal cortex outnumbered the other two levels in the common map (Table 4; For activation thresholds at p<0.001, LH: χ2 (2,2503) = 1936.16, p<0.001; RH: χ2 (2,1334) = 710.37, p<0.001; For thresholds at p<0.01, LH: χ2 (2,4682) = 2491.73, p<0.001; RH: χ2 (2,3749) = 1692.33, p<0.001). These results highlight that the differentiation of action classes was greatest in the parietal cortex, whereas the commonality was most evident in the occipito-temporal cortex.

**Table 4.** Comparison of the three levels of the AON in terms of the number of voxels that are active around the local maxima in specific maps and common maps with thresholds p<0.001 and p<0.01

|  | Specific Map for indirect communication |  | Common map for all three actions |  |
| --- | --- | --- | --- | --- |
|  | Left | Right | Left | Right |
| **$p < 0.001$** |  |  |  |  |
| Occipito-temporal | 24 | 52 | **1874** | **767** |
| Parietal | **362** | **364** | 90 | 120 |
| Premotor | 153 | 31 | - | - |
| **$p < 0.01$** |  |  |  |  |
| Occipito-temporal | 142 | 588 | **2715** | **1552** |
| Parietal | **929** | **1223** | 476 | 85 |
| Premotor | 397 | 301 | 23 | - |

#### 3.1.3 Activity Profiles of Action Classes

After the whole-brain mapping of regions involved (Section 3.1.1), specifically involved and commonly involved (See Section 3.1.2) in the observation of indirect communication, direct communication, and manipulation actions, the next step in the univariate analysis was to investigate how brain activity for the three action classes differed from one another within the pre-defined regions of interests at the three levels of the AON. For this purpose, we computed the activity profiles of the three action classes (videos) as well as their static and dynamic controls by computing the mean percent signal change in MR signal within each ROI (See Figure 5 for occipito-temporal and premotor ROIs, and Figure 6 for parietal ROIs).

**Figure 5.**
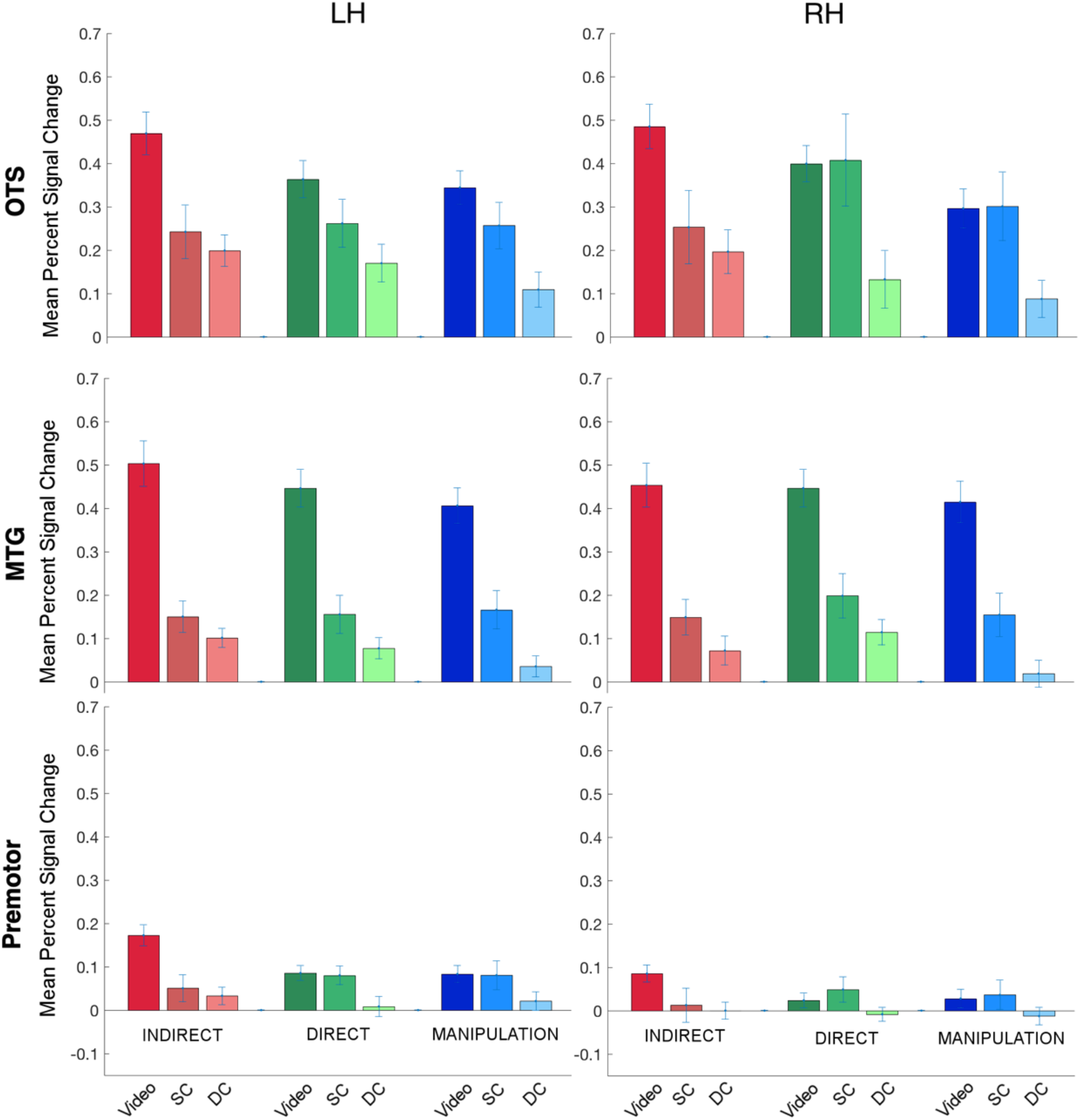
Activity profiles of the three action classes and their static and dynamic controls in ROIs in the occipito-temporal and premotor cortex. Video refers to the action stimulus. SC – Static control, DC – Dynamic control, LH – Left hemisphere, RH – Right hemisphere. Error bars show the standard error. (A) Left OTS and right OTS. (B) Left MTG and right MTG. (C) Left premotor and right premotor.

**Figure 6.**
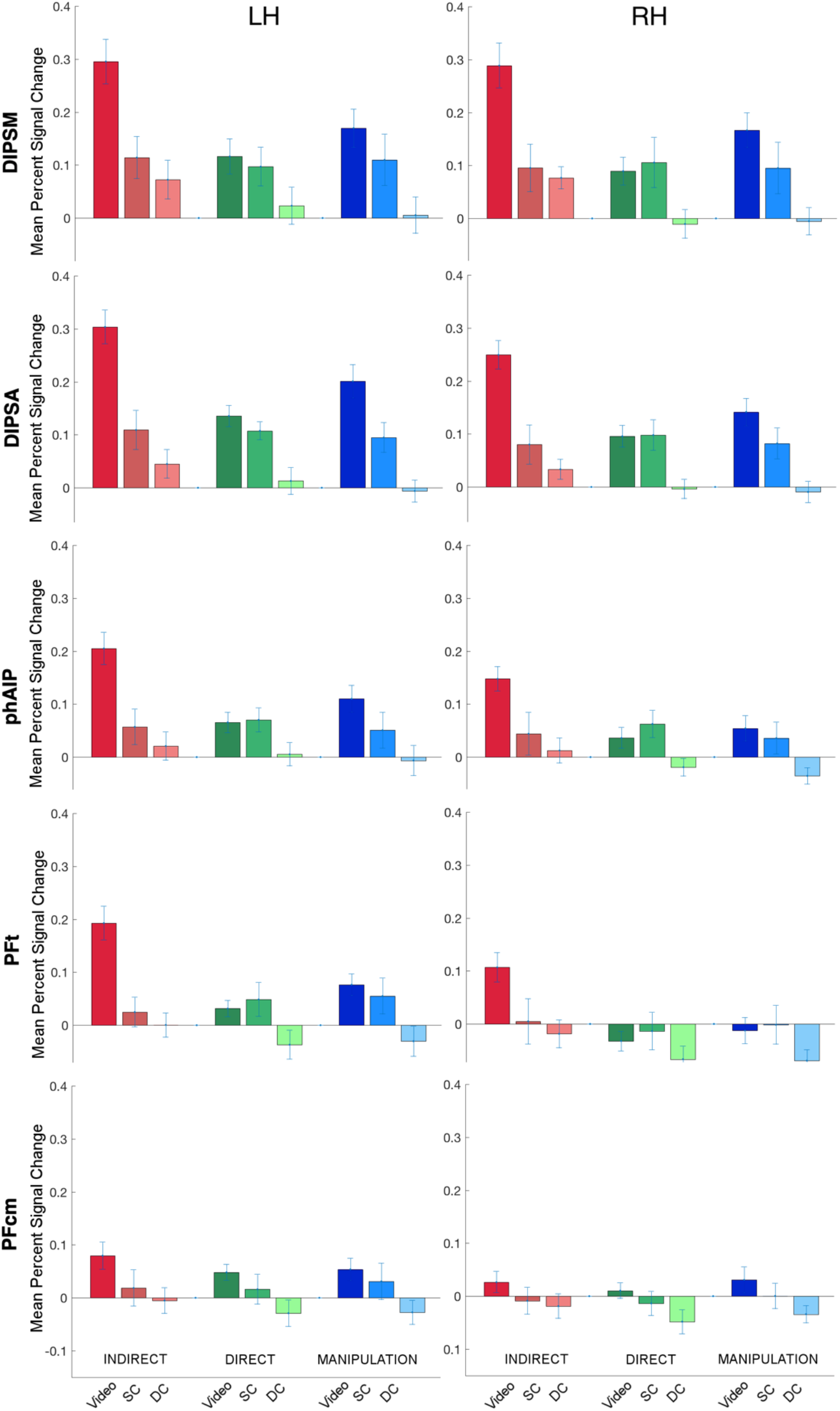
Activity profiles of the three action classes and their static and dynamic controls in ROIs in the parietal cortex. Video refers to the action stimulus. SC – Static control, DC – Dynamic control, LH – Left hemisphere, RH – Right hemisphere. Error bars show the standard error. (A) Left DIPSM and right DIPSM. (B) Left DIPSA and right DIPSA. (C) Left phAIP and right phAIP. (D) Left PFt and right PFt. (E) Left PFcm and right PFcm.

Overall, the mean percent signal change for indirect communication was greater than that of direct communication and manipulation for video presentation in all but a few ROIs. On the other hand, the relative activity of direct communication and manipulation varied across ROIs: the activity for direct communication was slightly higher than for manipulation in occipito-temporal areas, whereas it was lower in the parietal areas.

When we compared the activities for the videos with the static and dynamic controls, we found that the activity for video presentation was generally greater than that of the static and dynamic controls. Static controls generally induced stronger activity than did dynamic controls.

Comparing the hemispheres, left hemisphere displays higher activity overall than the right hemisphere, especially for direct communication and manipulation.

In order to examine the differences between conditions statistically, we performed an 8 (ROI) x 2 (Hemisphere) x 3 (Action class) x 3 (Presentation mode) mixed ANOVA including all variables of interest. This omnibus ANOVA showed a main effect of all four factors, ROI, Hemisphere, Action class, and Presentation mode. In addition, the two-way interactions ROI x Presentation mode, Action class x Presentation mode, ROI x Action class, the three-way interactions ROI x Action class x Presentation mode, ROI x Hemisphere x Action class, and the four-way interaction ROI x Hemisphere x Action class x Presentation mode were all significant. (See Table 5 for F-values and p-values of the main effects and the interactions).

**Table 5.** Main effects and interaction effects of the mixed ANOVA 8 (ROI) x 2 (Hemisphere) x 3 (Action class) x 3 (Presentation mode) on univariate brain activity (Significant effects are marked in bold in the first column)

|  | **F-value** | **p-value** |
| --- | --- | --- |
| **ROI** | $F_{7,357} = 56.64$ | **p<0.001** |
| **Hemisphere** | $F_{1,51} = 5.27$ | **p<0.05** |
| **Action class** | $F_{2,102} = 6.16$ | **p<0.005** |
| **Presentation mode** | $F_{2,51} = 21.95$ | **p<0.001** |
| ROI x Hemisphere | $F_{7,357} = 1.86$ | p=0.08 |
| **ROI x Action class** | $F_{14,714} = 6.70$ | **p<0.001** |
| **ROI x Presentation Mode** | $F_{14,357} = 7.69$ | **p<0.001** |
| Hemisphere x Action class | $F_{2,102} = 2.19$ | p=0.12 |
| Hemisphere x Presentation mode | $F_{2,51} = 0.98$ | p=0.38 |
| **Action class x Presentation mode** | $F_{4,102} = 4.49$ | **p<0.005** |
| **ROI x Hemisphere x Action class** | $F_{14,714} = 2.56$ | **p<0.005** |
| **ROI x Action class x Presentation mode** | $F_{28,714} = 2.50$ | **p<0.001** |
| ROI x Hemisphere x Presentation mode | $F_{14,357} = 0.45$ | p=0.96 |
| Hemisphere x Action class x Presentation mode | $F_{4,102} = 0.75$ | p=0.56 |
| **ROI x Hemisphere x Action class x Presentation mode** | $F_{28,714} = 1.93$ | **p<0.005** |

Pair-wise comparisons between ROIs showed interesting patterns. Overall, some anatomically nearby regions (e.g. OTS and MTG, DIPSM and DIPSA, PFt and PFcm) did not differ although they differed from the remaining of the ROIs (Figures 5–6). More specifically, MTG was significantly different from all other ROIs except OTS (p<0.001); OTS was significantly different from all ROIs except MTG (p<0.001); DIPSA was significantly different from all ROIs except DIPSM (p<0.001); DIPSM was significantly different from all ROIs except DIPSA (p<0.05); phAIP was significantly different from all ROIs except Premotor (p<0.05); PFt was significantly different from all ROIs except PFcm (p<0.05); PFcm was significantly different from all ROIs except PFt (p<0.05); and Premotor was significantly different from all ROIs except phAIP (p<0.05). Thus, in general OTC and PMC level differed from the parietal ROIs, which themselves showed some internal differences.

Pair-wise comparisons between the hemispheres showed that activity in the left hemisphere was significantly higher than in the right hemisphere (p<0.05).

On the other hand, pair-wise comparisons between the action classes showed that activity for indirect communication significantly exceeded that of direct communication and manipulation (p<0.05). Direct communication and manipulation did not significantly differ (Figures 5–6).

When we compared the presentation modes, we found that the video presentation elicited significantly higher activity than the static and dynamic controls (p<0.01). In addition, the static controls were significantly different from the dynamic controls (p<0.05) (Figures 5–6).

A further important pattern, reflecting the ROI x action class x presentation mode interaction, was that that the video conditions of the action classes differed more in parietal areas (DIPSM, DIPSA, phAIP, PFt), indirect communication being strongest and direct communication the weakest, compared to the other two levels, occipito-temporal (MTG) and premotor areas (Figures 5–6). The only exception to this pattern in parietal cortex was PFcm in which the action classes did not differ from each other for the video condition. On the other hand, there were no such differences between the action classes for static controls or dynamic controls.

#### 3.1.4 Activity Profiles of Exemplars

Observing a specific area for indirect communication action class in PFt motivated us to examine the activity profile of the exemplars. Our hypothesis was that if an area is specific for a particular action class, it should differentially respond to its own exemplars, but not to the exemplars of other action classes. On other words, we hypothesized that the activity profile of indirect communication exemplars would be significantly different from each other in PFt, but the activity profile of the manipulation exemplars or direct communication exemplars would not be different from each other in PFt. Furthermore, we hypothesized that this pattern of results would exist only in a region which specifically processes a particular action, i.e. PFt in the present study. To this end, we ran a one-way ANOVA in each parietal ROI separately (PFt, phAIP, PFcm, DIPSM, DIPSA) with factor as *exemplars* of all action classes (12 levels are drawing, erasing, sculpting, writing (indirect communication exemplars), clapping, right here, no sign, waving (direct communication exemplars), dragging, dropping, grasping, pushing (manipulation exemplars). Our results showed that there was a main effect of exemplars in all ROIs except PFcm (PFt: F(3.96,102.84) = 13.80, p<0.001; phAIP: F(4.64,120.66) = 10.89, p<0.001; F(4.43,115.26) = 11.84,p<0.001; DIPSM: F(4.66,121.14) = 19.92, p<0.001; PFcm: F(4.6,119.46) = 1.77, p=0.131). However, posthoc pair-wise tests revealed that only PFt (Figure 7) showed a significant difference between the exemplars of the indirect communication class, and not between the exemplars of the other two action classes (Figure 7) (writing vs. drawing: p=0.017; writing vs. erasing: p=0.001; writing vs. sculpting: p=0.000; drawing vs. sculpting: p=0.015; all other differences within the exemplars of a given action class: p>0.05). These difference between exemplars were specific for the video conditions (Figure 7). The main effects for the other ROIs were driven by the differences between the exemplars of different action classes, not within an action class.

**Figure 7.**
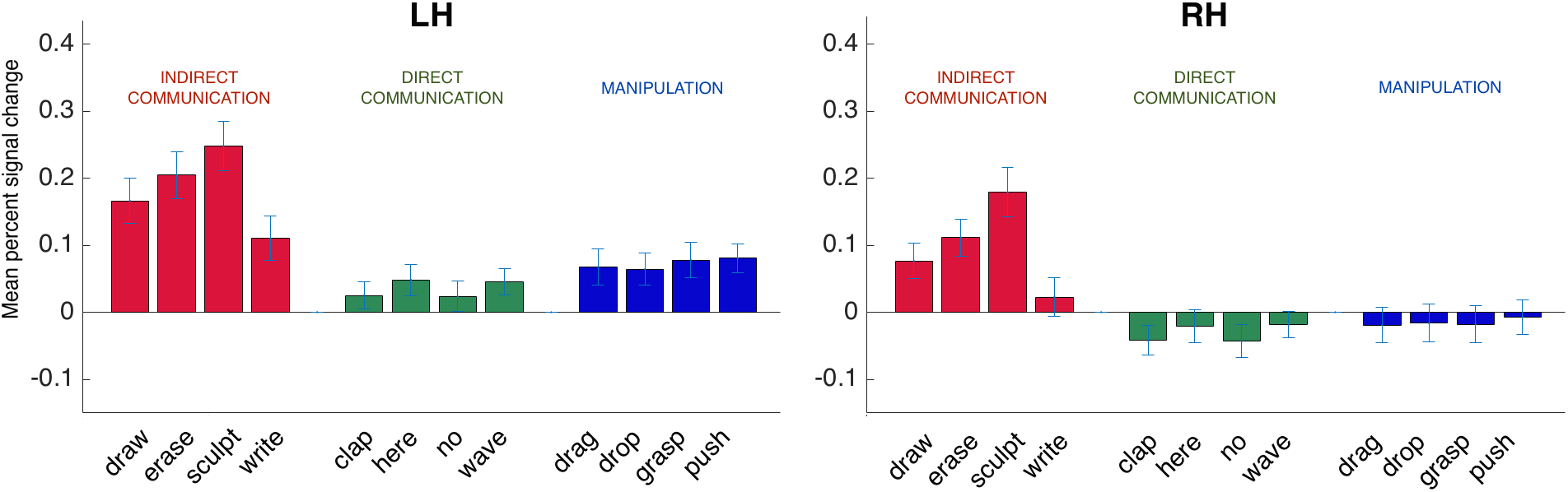
Activity profiles of the exemplars for all action classes in PFt. LH – Left hemisphere, RH – Right hemisphere. Error bars show the standard error.

### 3.2 MVPA: 3-way Pattern Classification across Action Classes

In order to investigate how well the three levels of the AON discriminated the indirect communication, direct communication, and manipulation actions, we applied a 3-way pattern classification to the neural activity patterns elicited by the video presentation of the three action classes in each ROI. We further investigated using the control conditions, whether the form or motion components of the observed actions could be sufficient to discriminate between the action classes. For this, we performed the 3-way pattern classification to the neural activity patterns both for the static controls (form information) and dynamic controls (motion information) of the three action classes. One should be cautious in interpreting the classification results for each presentation mode on its own, as videos, static frames and even the dynamic control videos are complex stimuli consisting of many components. For example, the static frames do not include the static hand alone but also the actor, a table, and a conspecific or a lamp on the left depending on the action class. Even the dynamic controls include the local motion related to the hand action, but at least for manipulation also the motion of the object resulting from the action.

Overall, the actions were discriminated significantly above chance in all ROIs for videos (chance 41%) as well as the static and dynamic controls (chance 44%, Figure 8). However, the discrimination patterns for videos and the controls differed. When we compared prediction accuracies between the videos and the controls, we found that parietal areas (including DIPSM, DIPSA, phAIP) showed higher prediction accuracies for videos compared to the static and dynamic controls, whereas occipito-temporal (left OTS) and premotor (left) ROIs showed higher prediction accuracies for the static and dynamic controls compared to videos. An 8 (ROI) x 2 (Hemisphere) x 3 (Presentation mode) ANOVA on the 3-way prediction accuracies showed a main effect of ROI, and a main effect of Presentation mode, as well as an interaction effect of ROI x Presentation mode (See Table 6 for the F-values and p-values). Pairwise comparisons between the presentation modes showed that the action class discrimination with the video presentation was significantly greater than the static control (p<0.001) but not the dynamic control, and that the dynamic control was significantly greater than the static control (p<0.01).

**Table 6.** Main effects and interaction effects of the ANOVA 8 (ROI) x 2 (Hemisphere) x 3 (Presentation mode) on 3-way prediction accuracy (MVPA) (Significant effects are marked in bold in the first column)

|  | **F-value** | **p-value** |
| --- | --- | --- |
| **ROI** | $F_{7,182} = 3.52$ | **p<0.01** |
| Hemisphere | $F_{1,26} = 2.83$ | p=0.10 |
| **Presentation mode** | $F_{2,52} = 14.78$ | **p<0.001** |
| ROI x Hemisphere | $F_{7,182} = 1.30$ | p=0.25 |
| **ROI x Presentation Mode** | $F_{14,364} = 6.97$ | **p<0.001** |
| Hemisphere x Presentation Mode | $F_{2,52} = 1.23$ | p=0.30 |
| ROI x Hemisphere x Presentation Mode | $F_{14,364} = 0.89$ | p=0.57 |

**Figure 8.**
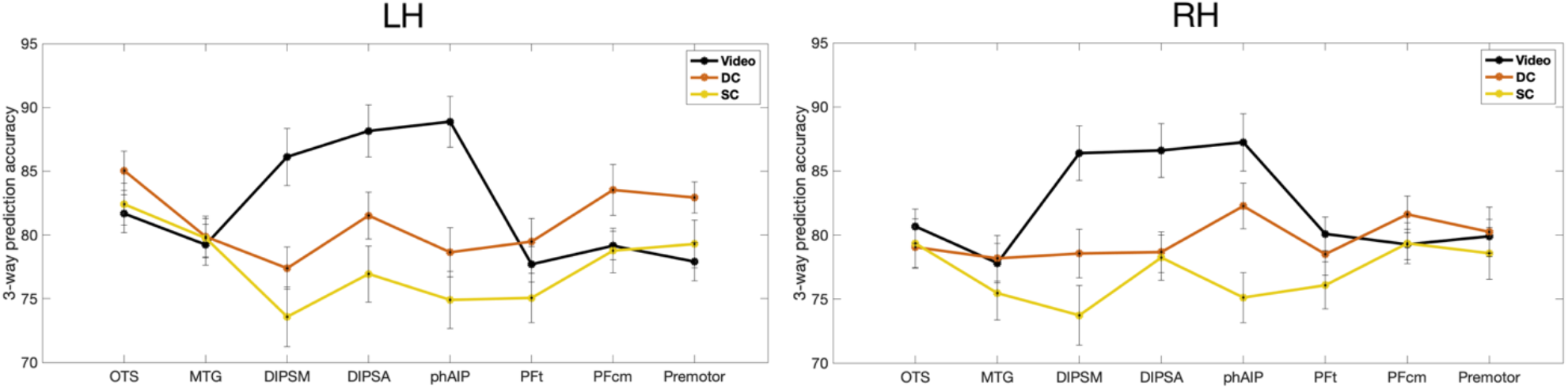
The mean prediction accuracy of between-action class classifications (3-way MVPA between Indirect, Direct, and Manipulation) for video, static control (SC), and dynamic control (DC) conditions in all levels of the Action Observation Network across 27 subjects. The results of left hemisphere are presented in the left panel and those of the right hemisphere in the right panel. The error bars show the standard error.

Next, we examined the discrimination with each presentation mode separately to further examine the variability across the three levels of the AON. For videos, the discrimination between action classes was much stronger in parietal cortex than in occipito-temporal and premotor cortex One-way ANOVA (ROI) of the 3-way prediction accuracy for videos showed a main effect of ROI in both hemispheres (Left: F(7,182) = 10.40, p<0.001; Right: F(7,182) = 7.38, p<0.001). Pair-wise differences between ROIs showed clear distinctions between the parietal cortex (especially DIPSA and phAIP) and the other two levels in the left hemisphere: DIPSA and phAIP discriminated the action classes significantly better than MTG and Premotor (p<0.05). Furthermore, there were several differences between parietal ROIs as well: DIPSA and phAIP discriminated significantly better than PFt and PFcm (p<0.01).

As expected from the interaction between ROIs and presentation mode, the discrimination between the action classes for the action videos was significantly better than static and dynamic controls in parietal regions (DIPSM: video vs. static control, t=6.86, p<0.001; video vs. dynamic control, t=4.51, p=0.002; DIPSA: video vs. static control, t=5.33, p<0.001; video vs. dynamic control, t=3.97, p=0.024; phAIP: video vs. static control, t=7.11, p<0.001; video vs. dynamic control, t=4.15, p=0.012). On the other hand, the discrimination between the action classes for static and dynamic controls, unlike the video presentation, was better in occipito-temporal than in parietal cortex: One-way ANOVA (ROI) of the 3-way prediction accuracy for both controls showed a main effect of ROI only in the left hemisphere (Static controls: F(7,182) = 2.59, p<0.05; Dynamic control: F(7,182) = 2.47, p<0.05), and pair-wise differences showed that for both controls OTS discriminated the action classes significantly better than DIPSM (p<0.05).

### 3.3 RSA

In order to investigate the neural representational distances between the three action classes, indirect communication, direct communication, and manipulation, and how they changed across the three levels of the AON, we employed representational similarity analysis. We computed the representational similarity matrix (RSM) for each ROI, and then applied hierarchical clustering and multidimensional scaling (MDS) on the dissimilarity matrices to visualize the distances between action classes and their exemplars for videos, static controls and dynamic controls.

For videos, the RSM for each ROI showed clear distinctions between the three action classes: the neural representations for the exemplars of each class were more similar to each other compared to exemplars of the other two classes. The RSMs of example ROIs from each level of the AON were shown in Figure 9A: MTG from left occipito-temporal cortex, phAIP from left parietal cortex, and left premotor cortex (other ROIs were not shown due to space limitations). The hierarchical clustering (Figure 9B) and MDS (Figure 9C) applied to the dissimilarity matrices (1-similarity matrix) showed this pattern more clearly wherein the exemplars of each action class were more closely clustered to each other compared to the exemplars of the other two action classes. In addition, hierarchical clustering also showed that the overall representational geometry was consistent across the three levels, in which the neural representation for manipulation and direct communication were more similar to each other compared to the indirect communication (See Figure 9B for the clustering structure in the dendrograms of the example ROIs). The only exception to this pattern was observed in left PFt in which the neural representations for direct and indirect communication were closer to one another compared to manipulation.

**Figure 9.**
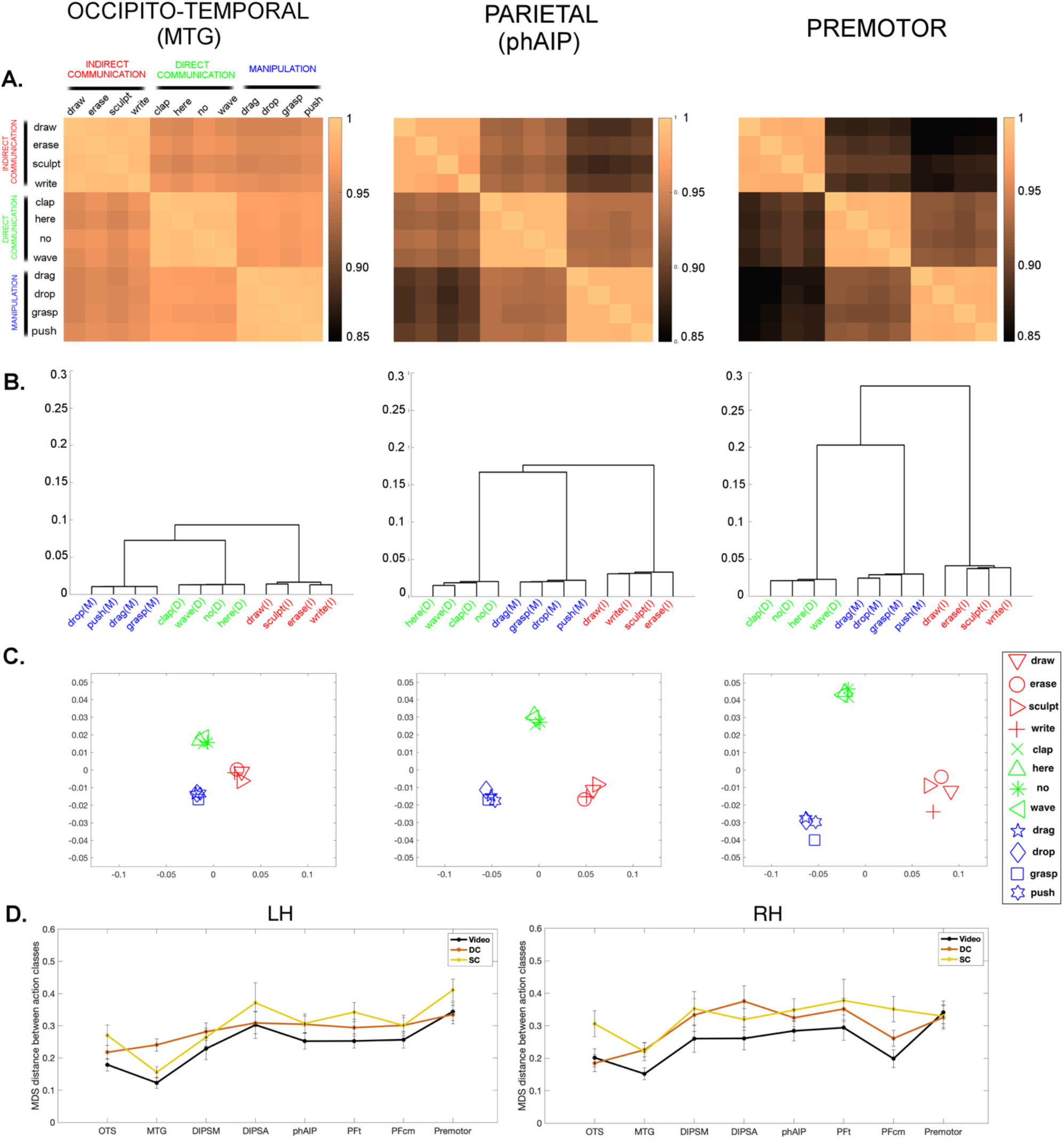
Representational similarity of action classes at the three levels of the Action Observation Network: A-C similarity for video condition in sample ROIs from each level: Left MTG from occipito-temporal, left phAIP from parietal, and left premotor from premotor cortex; D: similarity for all ROI and all presentation modes. (A) Representational similarity matrices for action videos in MTG, phAIP, and premotor. Similarity metric was correlation. Color bar shows the correlation strength. (B) Dendrograms obtained by applying hierarchical clustering to the dissimilarity matrices (1-similarity matrix, from panel A) (Left panel MTG, middle panel phAIP, right panel premotor). Each action class was marked with a different color following the conventions in Figure 5–6: Red shows the exemplars of Indirect Communication, green shows the exemplars of Direct Communication, and blue shows the exemplars of Manipulation. (C) Distance plots obtained by applying multidimensional scaling (MDS) to the dissimilarity matrices (Left panel MTG, middle panel phAIP, right panel premotor). (D) Sum of pair-wise distances between the three action classes for videos, static controls and dynamic controls on the MDS as in Panel (C) across all ROIs (LH: left hemisphere, RH: right hemisphere).

Although the neural representational geometry was consistent across the three levels of the AON, the representational distance between the action classes changed across the levels. Overall, the representational distance between the action classes was greatest in the premotor cortex, followed by the parietal cortex and occipito-temporal cortex as evidenced for the example ROIs by the height of the dendrograms in Figure 9B and geometric total distance in the MDS plots in Figure 9C.

In order to generalize these observations for the video conditions, and to clearly see the neural distances between each action class pair and how they changed across the three levels of the AON (similar to our approach in pattern classification, see Section 3.2.1.2), we plotted the pair-wise MDS distances between action classes, averaging over exemplars within each action class. The results were shown in Figure 9D (open circles). Overall, there was a quite consistent pattern in which distances between action pairs were greater in the premotor cortex followed by parietal, and by occipito-temporal cortex for videos. A 2 (Hemisphere) x 8 (ROI) ANOVA on the between-class differences confirmed these results. There was a main effect of ROI (F(7,182) = 7.88, p<0.001). Posthoc tests revealed that there were significant differences between MTG (occipito-temporal) and all other ROIs (MTG vs. DIPSM: p=0.04; MTG vs. DIPSA: p=0.006; MTG vs. phAIP: p=0.001; MTG vs. PFt: p=0.000; MTG vs. PFcm: p=0.083 (marginal); MTG vs. Premotor: p=0.000), as well as differences between OTS and Premotor (p=0.000), and DIPSM and Premotor (p=0.031). All other pair-wise differences were not significant (p>0.05). However, figure 9D shows that these changes in distance across ROI were very similar for the two control conditions. Indeed, An omnibus 3-way ANOVA including the presentation mode as a factor as well −2 (Hemisphere) x 8 (ROI) x 3 (Presentation Mode)-showed that there was an effect of ROI (F(7,182)=8.05, p<0.001) and presentation mode (F(2,52)=8.80, p<0.001) but no interaction between the two (F(14,364)=1.40, p=0.15), suggesting that the change across ROIs is similar for the three presentation modes (videos, static controls, and dynamic controls).

## 4. DISCUSSION

Our results indicate that parietal cortex hosts dedicated regions for observing human-specific indirect communication actions. They also show that action classes are most sharply discriminated at the parietal level of the AON, suggesting action identity is coded at this level. Finally, our study also reports some caveats in the use of multivariate techniques, particularly with regard to classification of complex stimuli.

### 4.1 Functional organization of the human parietal cortex for observed actions

The present study contributes to the growing body of literature indicating that observing different classes of actions activates different anatomical regions in human parietal cortex (Abdollahi et al 2013, Ferri et al 2015, Corbo and Orban, 2017). Observing indirect communication, a class of uniquely human actions, exclusively activated PFt and anterior phAIP as opposed to manipulation and direct communication. These results suggest that expansion of human IPL during evolution (Van Essen and Dierker, 2007) may have enabled the representation of action classes specific to humans. The specific PFt activation during observation of indirect communication actions may be accounted for by several sensorimotor differences between action classes, which assumes that observing and planning actions of a given class overlap in PPC (Ferri et al 2015). First of all, there is an overlap between the PFt activation and aSMG, an area involved in observing tool actions (Peeters et al 2009). Despite the absence of tools, indirect communication actions may have inherent tool-like kinematics (Peeters et al 2013) since the hand was used in a rigid manner imitating a tool (e.g. the index finger in drawing and writing). In fact, a previous study (Kroliczak and Frey, 2009) contrasting tool actions with communicative gestures (similar to direct communication) reported activation of similar parietal regions. Second, indirect communication, unlike actions of the other classes, may require continuous monitoring of peri-personal space to know when to terminate the action (Ghafouri and Lestienne, 2000; Brozzoli et al, 2010; Patane et al, 2020). This differential association of sensorimotor with spatial processing may be one driving force for the specificity of PFt activation. Finally, PFt may maintain the “mind’s eye” of the trace to be left on the substrate in indirect communication actions, as has been postulated for planning artwork (Morris-Kay, 2009). The other classes do not require such a mental image: here external signals guide the transformation: size and orientation for manipulation (e.g. to know how to grasp, Culham and Valyear, 2006), or the height of the human target for direct communication (e.g. to know how much to raise the hand to wave).

Our results also qualify the role of phAIP whose involvement in observing manipulation is well established (Jastorff et al 2010; Abdollahi et al, 2013; Ferri et al 2015; 2016; Corbo and Orban, 2017; Orban et al, 2019; Tyson et al, 2020). Bilateral phAIP was involved in manipulation and indirect communication actions, which correspond to relatively similar sensorimotor transformations as the hand interacts with either an object (with the goal of displacing or deforming) or a substrate (with the goal of transforming). In other words, indirect communication involves modifying the substrate, which is similar to manipulating objects – you cannot move the substrate but you deform it, and the sensory input for its planning is the 3Dshape of the substrate and the 3D texture which are also used in manipulation. On the other hand, the absence of phAIP involvement in observing direct communication is reminiscent of the distinction between observing intransitive and transitive actions, the former failing to activate the rostral part of IPS relative to seeing a moving dot (Jonas et al 2007). Instead, direct communication actions activated area PFcm (at the parietal level), an area which was previously shown to be activated by vocal communication actions (e.g. speaking, Corbo and Orban, 2017).

### 4.2 Comparison of the three levels of the AON

Our study reveals that the parietal level segregates the observed action classes more distinctly than the two other levels. Indeed, the specific map included larger regions at the parietal level than at the two other, while the common map was more extensive in the OTC than at the other two (Table 4). The parietal level also exhibited the greatest univariate separation (Figures 5–6), and best discrimination between the videos of the three action classes (Figure 8), an effect specific for the videos compared to controls.

Together, these findings shed light on the role of each level in action observation. This and previous studies suggest that the highest level of observed action identity, action class, is processed in anatomically segregated PPC regions, in all likelihood because action observation shared neural structures with action planning (Ferri et al 2015). These structures, the “intentional maps” (Anderson and Buneo, 2002), implement the sensorimotor transformations used in the definition of action classes. That many of the class-specific PPC regions are located in rostral PPC is consistent with the findings of Roth and Zohary (2015) that position invariance increases along a caudo-rostral gradient in PPC.

If the occipito-temporal and premotor levels of the AON do not process observed action identity what, then, is their contribution? Given that the OTC regions are shared by many action classes, they may process factors common to these many different actions such as the actor (Orban 2015, Cross et al 2009b), or more abstract factors such as agency (Sperduti et al 2011). There is also evidence that OTC regions code some abstract aspects of observed actions such as category of target objects (Wurm and Lingnau, 2015), transitivity (Wurm et al, 2017), or sociality and interaction envelope (Tarhan and Konkle, 2020), which might result from using actions with man-made objects (e.g. tools, boxes) or complex scenes (e.g. using a knife in the kitchen), especially when presented as static stimuli. On the other hand, recent evidence suggests that the number of fingers used in grasping is encoded at the premotor level (Fabbri et al 2016). This is consistent with the suggestion that in action observation the premotor cortex processes the effector of the action, because premotor, and not parietal cortex exhibits a somatotopic organization (Buccino et al 2001, Jastorff et al 2010). This is supported by the lack of class segregation in the premotor cortex, because the effectors used in the three observed action classes were matched, and by the increase in representational distance between the *exemplars*, which differ in effectors, in premotor cortex compared to the other levels (Figure 9C).

### 4.3 Cross-method comparison between univariate analysis, MVPA, and RSA

In the present study, univariate analysis and multivariate techniques were complementary, but differences in the results provided us some caveats in using these techniques.

First, we note that the specification of the spatial extent of an ROI is crucial. In the present study, the lack of differences between action classes in PFcm with univariate analysis as opposed to the above-chance performance in the MVPA could be explained by the distinct activation patterns in the dorsal and ventral PFcm for manipulation and communication actions. Earlier research with manipulation actions showed activation in the relatively more dorsal part of PFcm (Ferri et al 2015) whereas earlier research with vocal communication actions showed activation in the relatively ventral part of PFcm (Corbo and Orban, 2017) (See Figure 4B). These findings suggest that communication and manipulation actions were represented by distinct population responses in PFcm and that the univariate analysis might have missed that pattern information by averaging across voxels, which may also be the case for some studies that reported lack of difference between manipulation and communicative actions at the parietal cortex (Montgomery et al 2007).

Second, the interpretation of MVPA results in fMRI experiments using complex stimuli such as natural images or videos may be difficult if control stimuli are not used. The videos as well as the static and dynamic controls showed above-chance performance. This raises the possibility that not only the videos, but also the controls, includes several features other than the action, some of them complex, which differed between classes and were apparently utilized by the decoding algorithms. This especially becomes a problem when the feature is linked to a single class, such as the conspecific on the left of the direct communication videos. Hence our results underscore the superiority of a classification of multiple items (and certainly more than two), and the desirability of decoding of control stimuli to verify the specificity of the classification, in line with caveats for interpreting fMRI MVPA results in terms of neuronal selectivity (Dubois et al, 2015).

The final point concerns the comparison of multivariate techniques, given the discrepancies between MVPA and RSA results: Although both methods showed separation between the three levels of the action observation network, the separation shown by MVPA was specific to video stimuli as opposed to the control stimuli whereas RSA separation was similar for all presentation modes. MVPA showed the greatest separation at the parietal level. This discrepancy suggests that although both techniques use multivariate data, the different metrics reveal distinct aspects of the data, with discriminability (MVPA) being a more discrete measure (above-chance or not), and distance (RSA) more continuous. Furthermore, MVPA can detect subtle biases in voxels across conditions (Kamitani and Tong, 2005) to which RSA may be less sensitive. Regardless, the fact that MVPA shows distinctions between the three levels of the action observation network adds further evidence that the three levels process different aspects of the observed actions.

## Acknowledgments

This study was supported by ERC AdG-2012 323606 (Parietal action) to Guy A. Orban. The authors would like to thank to Stefania Ferri and Daniele Corbo for their help in preparation of stimuli and data collection. The authors declare no competing financial interests.

